# Extending the Near Infrared Emission Range of Indium Phosphide Quantum Dots for Multiplexed *In Vivo* Imaging

**DOI:** 10.1101/2021.02.03.429632

**Authors:** Alexander M. Saeboe, Alexey Y. Nikiforov, Reyhaneh Toufanian, Joshua C. Kays, Margaret Chern, J. Paolo Casas, Keyi Han, Andrei Piryatinski, Dennis Jones, Allison M. Dennis

## Abstract

This report of the reddest emitting indium phosphide quantum dots (InP QDs) to date demonstrates tunable, near infrared (NIR) photoluminescence and fluorescence multiplexing in the first optical tissue window with a material that avoids toxic constituents. This synthesis overcomes the InP synthesis “growth bottleneck” and extends the emission peak of InP QDs deeper into the first optical tissue window using an inverted QD heterostructure. The ZnSe/InP/ZnS core/shell/shell structure is designed to produce emission from excitons with heavy holes confined in InP shells wrapped around larger-bandgap ZnSe cores and protected by a second shell of ZnS. The InP QDs exhibit InP shell thickness-dependent tunable emission with peaks ranging from 515 – 845 nm. The high absorptivity of InP leads to effective absorbance and photoexcitation of the QDs with UV, visible, and NIR wavelengths in particles with diameters of eight nanometers or less. These nanoparticles extend the range of tunable direct-bandgap emission from InP-based nanostructures, effectively overcoming a synthetic barrier that has prevented InP-based QDs from reaching their full potential as NIR imaging agents. Multiplexed lymph node imaging in a mouse model shows the potential of the NIR-emitting InP particles for *in vivo* imaging.

---

Fluorescence is an important biomedical imaging modality because of the relatively low cost of imaging equipment, nominal toxicity of non-ionizing radiation (*i.e*., light), potential for molecular imaging using target-specific contrast agents, and the prospect of multiplexed imaging using discretely colored fluorophores.^1, 2^ Molecules common in biological tissues including lipids, water, and hemoglobin both scatter and absorb light, limiting the ability of visible wavelengths to penetrate tissue structures.^3^ However, tissue-penetrating fluorescence imaging is feasible in the first optical tissue window (650-950 nm), where both light scattering and absorption are reduced compared to visible wavelengths.^4^ Semiconductor quantum dots (QDs) are inorganic nanoparticle emitters exhibiting size-dependent tunable emission peaks and broad, overlapping absorption peaks that are ideal for multiplexed imaging. While traditional cadmium selenide-based QDs are somewhat limited in their preclinical imaging applications because the CdSe bulk bandgap of 712 nm (1.74 eV) limits their emission range to visible wavelengths, the bandgap of indium phosphide at 925 nm (1.34 eV) promises tunability and multiplexing capacity throughout the NIR-I optical tissue window.^5, 6^ Cadmium-free QDs exhibit considerable promise for tissue-depth imaging both because of the advantages of the extended wavelength range and because of an interest in moving away from heavy metal-laden materials in the hopes of eliminating toxicity concerns.^7-11^

Recent advances in InP synthesis include the development of novel synthetic schemes that drive down the cost and environmental impact of InP synthesis while also improving photophysical performance.^12-15^ These Type I and Quasi-Type II InP QDs exhibit high quantum yields with reasonably narrow emission FWHMs (< 40 nm),^15, 16^ but do not emit throughout the first optical tissue window; reported emission peaks for colloidal InP are all below 750 nm, regardless of synthesis method.^12, 15-19^ Indium phosphide cores with cadmium sulfide or cadmium selenide shells emit at wavelengths >1000 nm, but the Type II band alignment of these structures coincides with longer photoluminescence lifetimes and lower quantum yields in addition to the unfortunate inclusion of cadmium.^20-23^ In order to address the shortcomings of InP tunability in the NIR, copper-doped InP and ternary systems such as copper indium sulfide have been developed.^24-26^ These systems exhibit dopant-based NIR emission, but this emission mechanism is thought to inherently limit the maximum brightness of the emitters and increase the emission peak width. ^24–29^ Our recent work describing CdSe shell emission demonstrated that lower energy/longer wavelength peak emission was feasible with confinement in the QD shell layer than with the same volume of CdSe in the traditional spherical QD morphology.^22, 23^ Given concerns that a “growth bottleneck” precludes synthesis of InP cores large enough to produce emission peaks spanning the entire NIR emission range theoretically possible for this material,^30^ we hypothesized that an InP shell structure may enable lower energy emission than we have observed with InP cores.

In traditional Type I QDs, both the electron and hole reside in the semiconductor core, the size and composition of which dictates emission energy/wavelength/color. In Quasi-Type-II heterostructures, one charge carrier (electron or hole) is confined, but band offsets in either the conduction or valence band are close enough in energy that the excited electron or hole is delocalized.^6^ In the prototypical case, *i.e*., CdSe/CdS, the electron wave function spreads from the core into the shell material, while the wave function of the heavy hole is confined to the core.^31^ In an *Inverse*-Type I semiconductor nanoparticle, a high bandgap core material is coated with a second semiconductor with a smaller bandgap and lower energy valence and conduction bands, confining the excited electron and the hole in the shell.^6^ Cadmium-based Inverse-Type I structures were briefly mentioned in the QD literature over 10 years ago,^32-34^ but yielded minimal subsequent interest until recently.^35^ A brief, singular report of thin-shelled, visible emitting ZnSe/InP/ZnS particles described them as Inverse-Type I structures.^36^ We synthesized the inverted structure with concerted methods promoting the deposition of thick, epitaxial semiconductor shells^37-39^ to generate cadmium-free NIR emission that pushes to longer wavelengths than reported thus far with Type I or Quasi-Type II InP core structures (**Figure 1, Figure S1**). These results highlight the advantages of non-conventional heterostructured QDs, while opening the door to further development of NIR-emitting InP nanoparticles.

**Figure 1.**
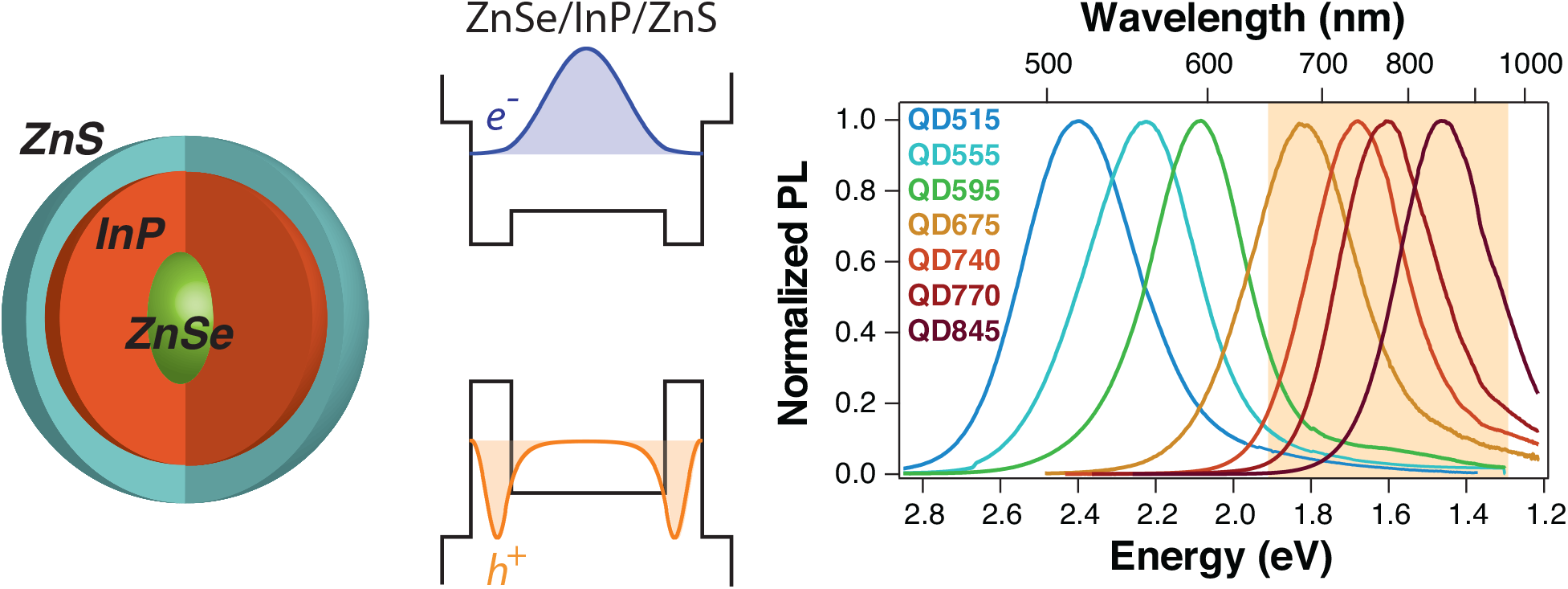
ZnSe/InP/ZnS core/shell/shell nanoparticles. The large-bandgap ZnSe core acts as a spacer around which the optically active InP shell is wrapped. The resulting InP QDs can emit at longer wavelengths than existing literature reports of InP QDs and are tunable with shell thickness. From left to right: schematic of the core/shell/shell structure design, diagram of the ZnSe/InP/ZnS bandgap alignments with corresponding electron and hole wavefunctions, and normalized photoluminescence (PL) spectra of QDs with increasing InP deposition. Note the electron and hole densities presented are for a 3.4 nm ZnSe core with a 0.75 nm InP shell; the ZnS capping shell was modeled as an infinite well.

The electronic structure of ZnSe/InP was calculated using a two-band effective mass model; the effect of the high bandgap ZnS cap (second shell) is negligible and was not accounted for in the calculations. The specific parameters used in the calculation are outlined in the SI. The result of the modeling shows that ZnSe/InP heterostructures with sufficiently thick InP shells are predicted to exhibit Inverted-Type I electron and hole localization to the shell (**Figures S2**), while particles with less than 3 nm thick InP shells, like those demonstrated previously,^36^ exhibit quasi-Type II band structure with the electron delocalized over the core and shell and the hole localized to the shell (see band diagram in **Figure 1, Figure S3**). In either case, the ‘heavy hole’ of InP localizes to the InP shell as soon as it is > 0.5 nm thick. This model predicts that the emission color of the ZnSe/InP structure can be tuned by changing the InP shell thickness. The bandgaps predicted by EMM indicate that both traditional^22, 23^ and inverted QD structures are theoretically capable of producing tunable emission throughout the first optical tissue window, but synthetic experience indicates that this has been technically challenging to achieve for InP core-based QDs, as these NIR emissions are not reported in the literature.

Inverted InP-based heterostructure reactions proceeded using hot-injection colloidal chemistry methods. First, zinc selenide cores were formed in a high-temperature, inert atmosphere precipitation reaction between zinc oleate and selenium complexed by diphenyl phosphine (Se:DPP), similar to previous reports.^40^ The resulting ZnSe cores exhibit a clear 1S absorption peak, indicating quantum confinement and reasonably narrow size distribution (**Figure S4**). The ZnSe cores were purified in a glovebox under argon *via* ethanol precipitation, then resuspended in octadecene (ODE) with 10% (v/v) oleylamine (OAm). Under argon atmosphere on a Schlenk line, indium and phosphorous precursors were added via a modified successive ionic layer adsorption and reaction (SILAR) approach.^41^ In a typical SILAR reaction, shell layers are added one ionic layer at a time through dropwise addition of the precursors followed by a high-temperature anneal (178°C; 1 or 2.5 hr) to ensure epitaxial crystal growth and reduce nucleation of the shell material. The indium and phosphorous precursors were 0.2 M indium oleate (1:3.3 In:oleic acid (OA)) in ODE and 0.2 M tris(trimethylsilyl)phosphine ((TMS)_3_P) in ODE, respectively. InP is highly susceptible to oxidation and these surface defects inhibit photoluminescence from the InP nanostructure,^42^ so a second, protective shell of zinc sulfide was added via SILAR using zinc oleate (0.2 M 1:3.3 Zn:OA) and 0.2 M sulfur dissolved in ODE to produce strongly emissive and chemically stable particles. The quantum yield (QY) of as-synthesized particles was sometimes further increased using a post-synthesis treatment process whereby cleaned particles were heated to 240°C for 3 hrs with 0.2 M zinc oleate in trioctylphosphine (TOP), producing zinc-terminated particles.

Up to 16 rounds of SILAR were used to deposit the InP immediately followed by 2-3 rounds of ZnS SILAR. As can be seen in the bright-field (BF) scanning transmission electron microscopy (STEM), the QDs gradually increase in size, indicating progressive growth of the InP layer with additional SILAR iterations (**Figure 2a Figures S5-S6**). Microwave plasma atomic emission spectroscopy (MP-AES; Agilent 4200MP-AES) of acid-digested samples yields the elemental composition of the core/shell/shell structures. Atomic concentrations in ppb were converted to molarity to calculate the relative abundance of the discrete elements for each sample. By combining the elemental ratios with the average particle size from STEM images and the assumption that the particles are concentric spheres comprising ZnSe/InP/ZnS, we estimated the relative core and shell dimensions for the QDs (**Figure 2b**). We assume that the relative volumes of the ZnSe core, InP shell, and ZnS shell are proportional to the molar ratios of Se, In, and Zn-minus-Se, respectively, adjusted for the crystal densities of each of the semiconductors. These results show that the InP shell grows progressively with an increased number of InP SILAR shelling iterations (**Figure 2b, Table 1**). The ZnS capping shell is ∼1 monolayer thick for each of the measured samples, consistent with the sub-quantitative reaction efficiency often seen with the Zn(OA)_2_-based shelling protocol.

**Table 1.**
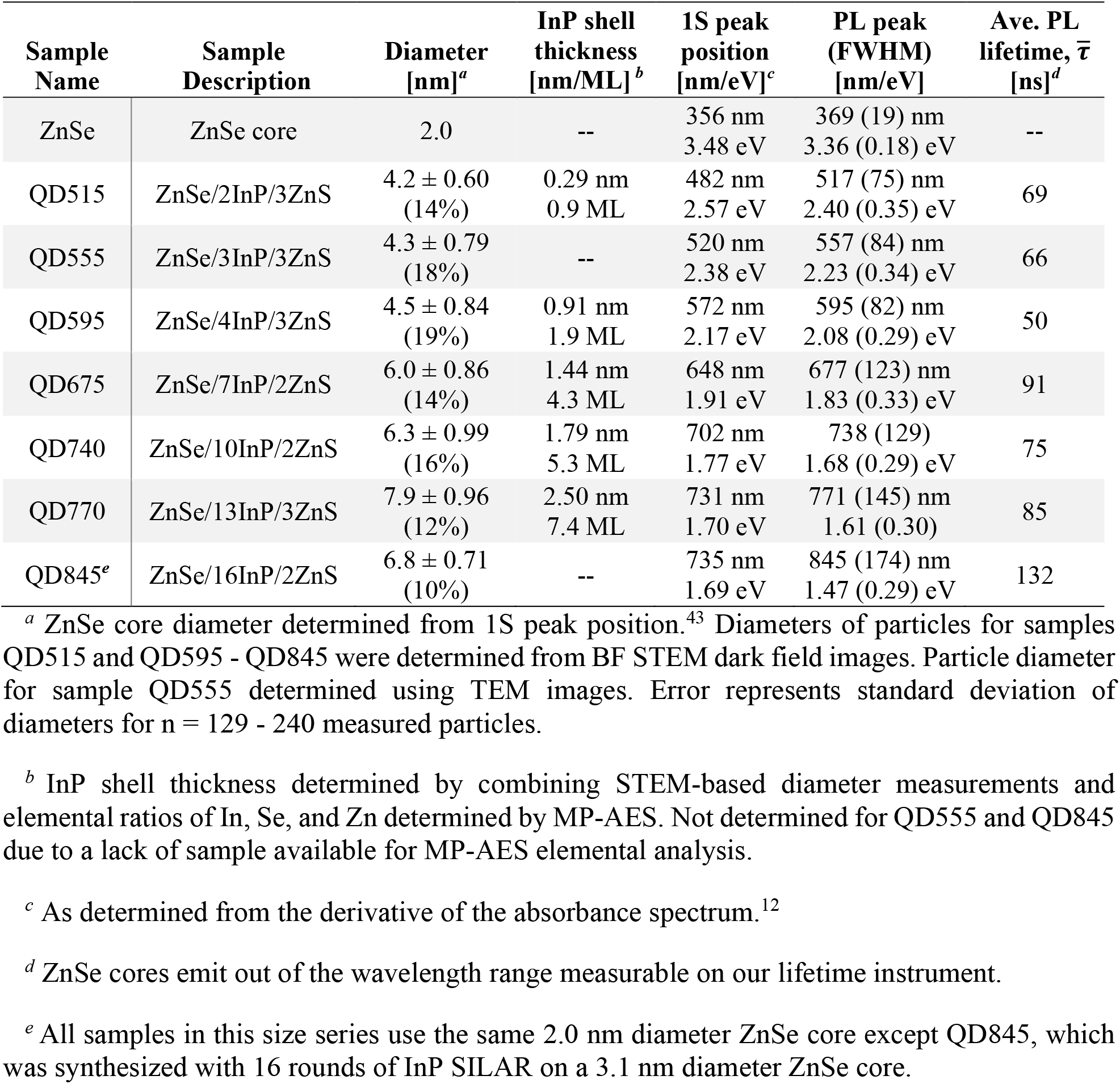
Summary of QD properties.

**Figure 2.**
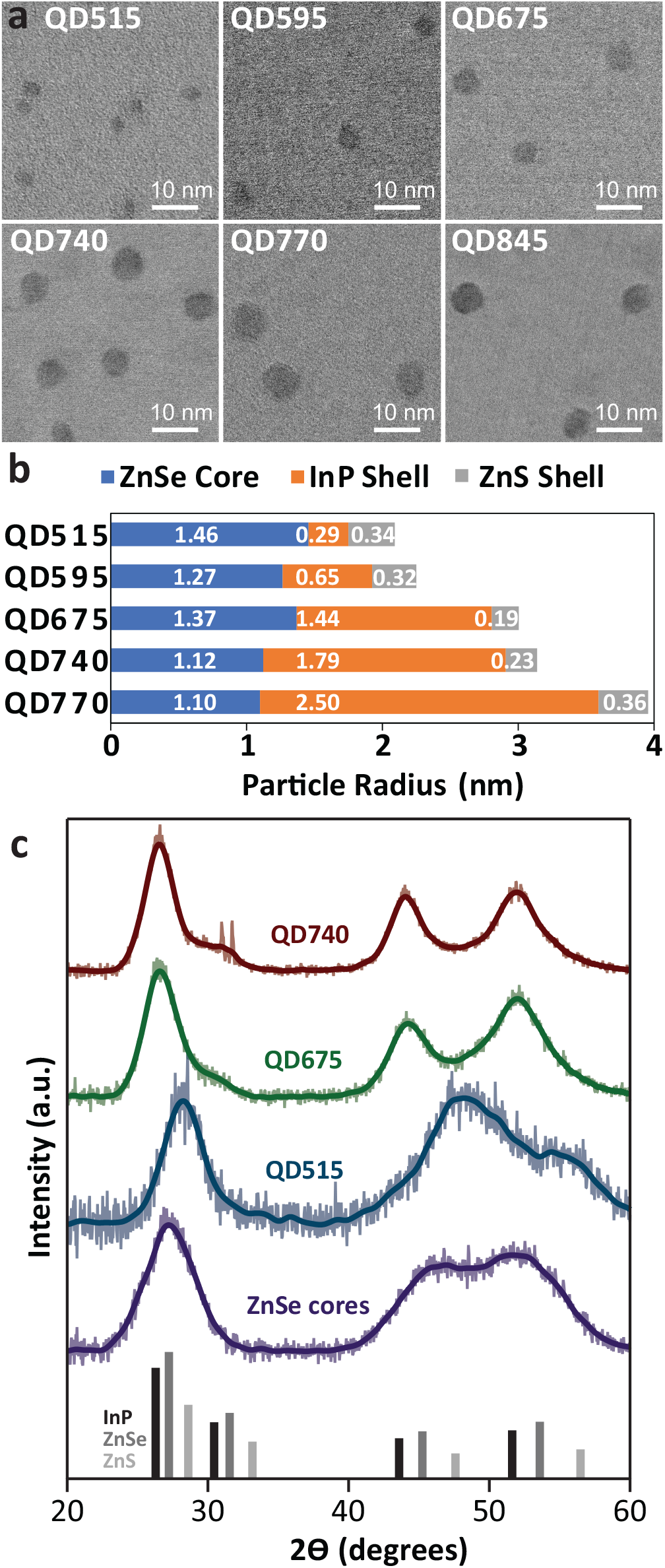
Materials characterization of QDs. (**a**) Bright-field (BF) STEM images of samples QD515, QD595, QD675, QD740, QD770, and QD845. (**b**) ZnSe/InP/ZnS core/shell/shell dimensions based on atomic analysis (MP-AES), assuming concentric spheres with total diameter equal to the average diameter determined by STEM. The sample is indicated on the left with the component dimension noted in nm on the bar section corresponding to the ZnSe radius (blue), InP shell (orange), and ZnS shell (grey). (**c**) X-ray diffraction (XRD) spectra of the ZnSe cores and several QDs, along with vertical lines representing reference InP, ZnSe, and ZnS peaks from the Crystallography Open Database (COD).

Precursor deposition and nanocrystal growth is also seen in the X-ray diffraction spectra of the materials (**Figure 2c**). The primary low angle diffraction peak of sample QD515, observed at 28.2°, is at a considerably higher angle than the ZnSe core peak, observed at 27.2°, and close to the ZnS peak position 28.5°. This shift towards a characteristic ZnS X-ray diffraction pattern suggests that the QD515 sample volume is dominated by ZnS. In contrast, samples QD675 and QD740, with primary low angle diffraction peaks at 26.6° and 26.5°, respectively, approach the characteristic InP diffraction peak at 26.3°, indicating that these QD largely comprise InP by volume. These sample diffraction peak shifts are consistent with the volume predominance expected for the samples, with a substantial ZnS contribution for samples with few InP SILAR cycles, and InP majorities for samples with larger numbers of InP SILAR cycles. Nanocrystal growth is also clearly observed based on a narrowing of the XRD peaks for samples with larger numbers of InP SILAR cycles. This narrowing is particularly evident in the 40-60° range: the two characteristic zincblende diffraction peaks of the ZnSe cores and QD515 in this region are sufficiently broad that the individual peaks are not readily resolvable, while samples QD675 and QD740 have two clearly resolvable peaks. Further growth of the nanocrystals is also supported by a reduction in full width half maximum (FWHM) of the primary low angle diffraction peak from 2.8° to 2.3° from QD675 to QD770, respectively. In addition to crystal size and composition, the XRD peak position and width could also be affected by strain effects; however, these strain effects are difficult to predict and therefore are not considered explicitly.^44, 45^

The presence of constituent elements in QD clusters was examined using energy dispersive spectrometry (EDS) mapping with Super X EDS detector on Tecnai Osiris TEM operated in STEM mode (**Figure 3**). EDS elemental mapping was performed on the samples before and after deposition of the outer ZnS shell. Since the electron beam excites X-rays from both ZnSe core and its outer InP and ZnS shells, the elemental maps show a relative elemental distribution averaged over the entire particle volume probed. The elements present in the mapped dots include zinc and selenium from the cores and indium and phosphorus from the inner shells with sulfur detected at 2.3 keV after addition of the outer ZnS shell. Also appearing in the EDS spectra are the energy lines corresponding to copper from the grid with supporting carbon membrane.

**Figure 3.**
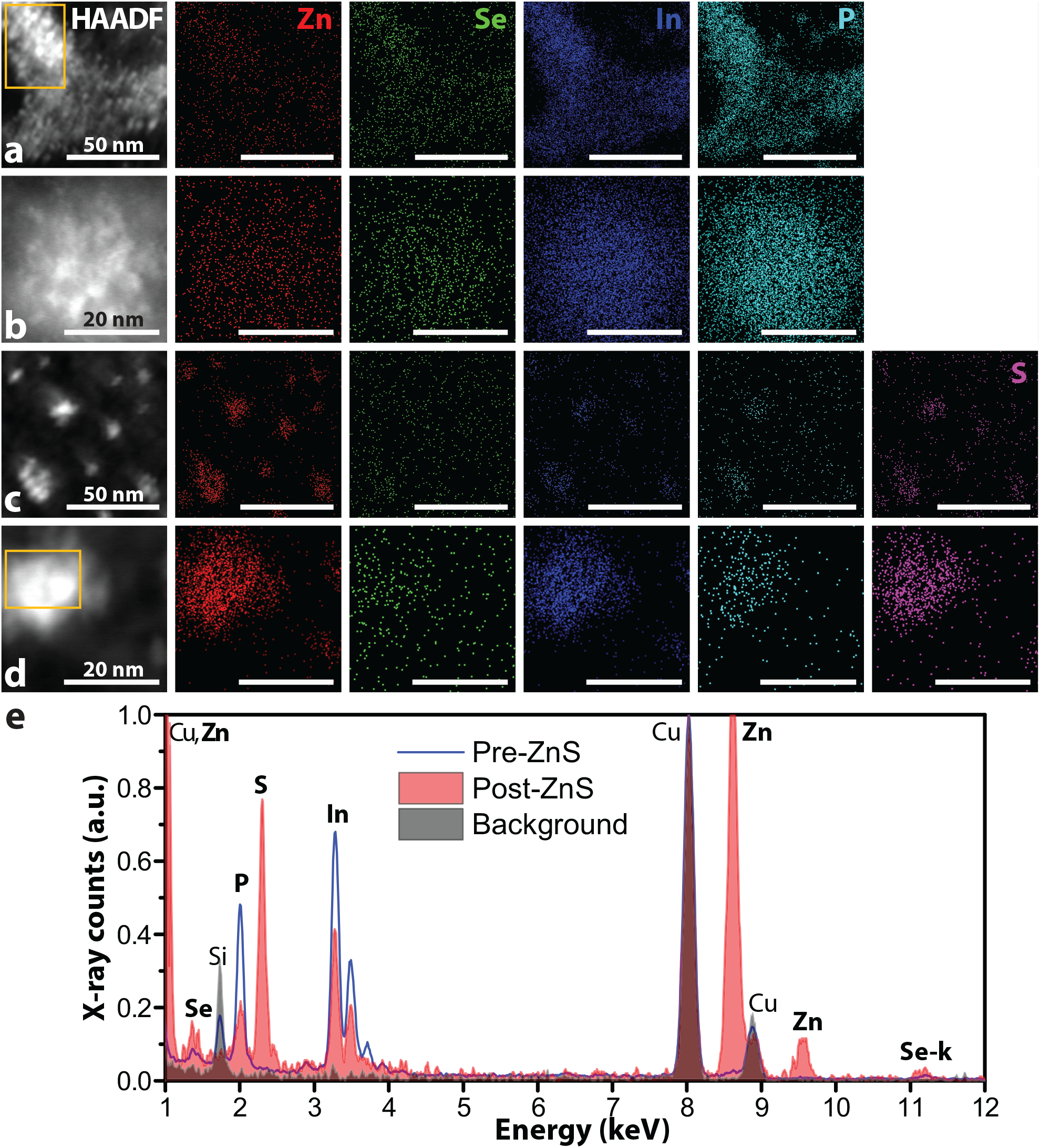
Elemental dispersive X-ray spectroscopy (EDS) Elemental maps showing the distribution of each element of interest alongside a reference high-angle annular dark field (HAADF) image indicating the spatial position of particle clusters; scale bars in the elemental maps are the same as the corresponding HAADF image. QD515 (**a**,**b**) before and (**c**,**d**) after ZnS capping. To generate the elemental maps, the X-ray K lines were used for Zn, P, and S, while X-ray L lines were used for In and Se. (**e**) EDS spectra collected from yellow inset regions in the reference HAADF images show the presence of elements in a cluster of QD. A representative background spectrum is included for reference; spectral intensities are normalized to the copper peak at 8 keV.

The QDs exhibit InP deposition-dependent quantum confinement and optical properties (**Figure 4**). InP QDs grown on the same ZnSe cores emit redder with additional InP shell depositions. For example, the synthesis of QD515, QD555, QD595, QD675, QD740, and QD770 all used the same 2.0 nm diameter ZnSe core combined with 2, 3, 4, 7, 10 and 13 rounds of In and P SILAR addition, respectively, and 2-3 rounds of ZnS addition. The InP shelling reaction was repeated with subtle changes to the reaction conditions on different batches of ZnSe cores, yielding similar tunability with InP addition (**Figure S7, Table S1**). Of these samples, only QD845, which was synthesized with 16 rounds of SILAR on a 3.1 nm diameter ZnSe core, is included together with the primary series to avoid redundancy.

**Figure 4.**
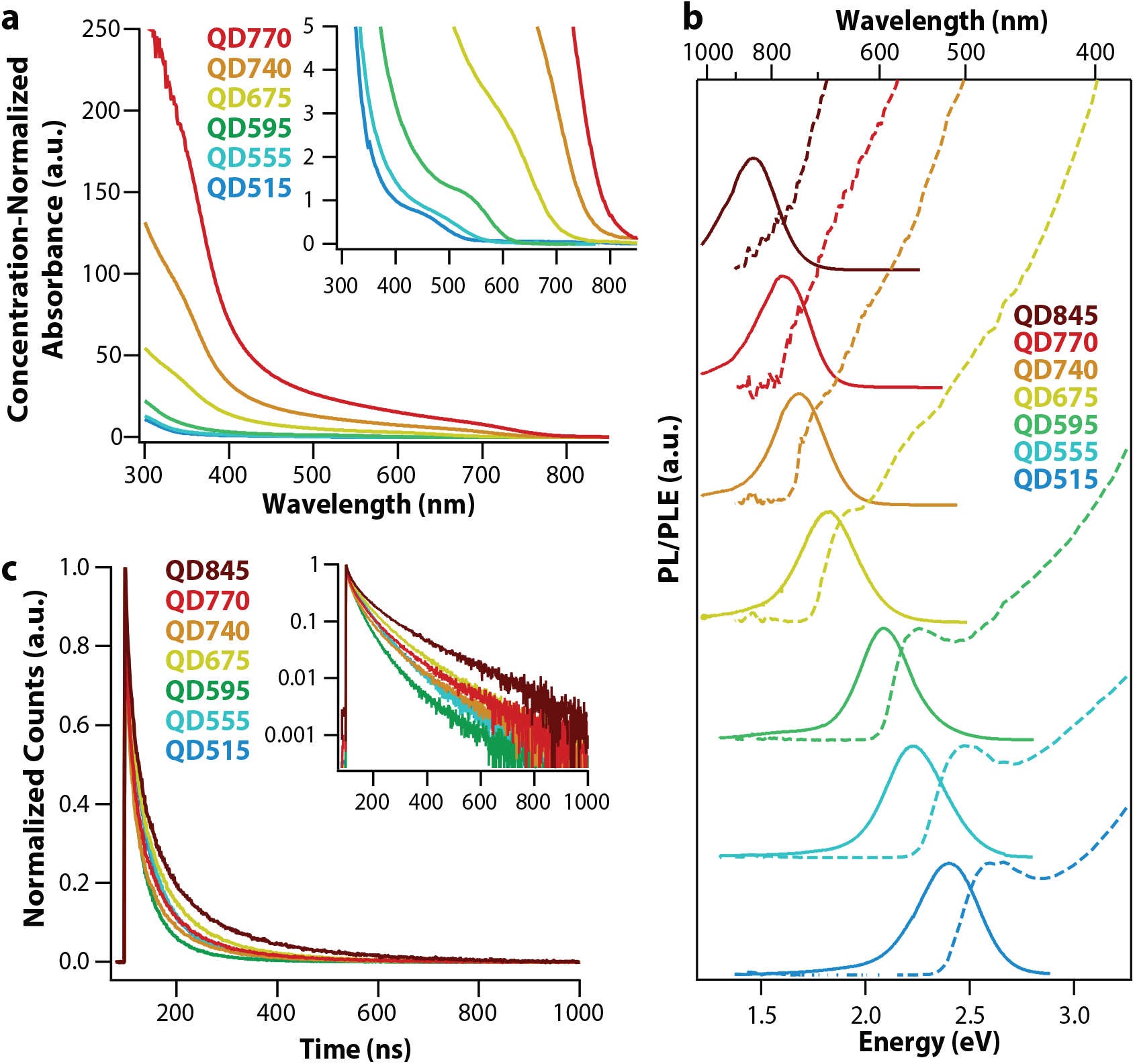
Optical properties of QDs. (**a**) Concentration-normalized absorbance was determined by measuring the absorption of carefully diluted reaction products (unwashed) from reactions loaded with identical amounts of the same ZnSe core and adjusted for reactant addition volumes. (**b**) Normalized photoluminescence and photoluminescence excitation spectra. (**c**) Photoluminescence lifetime following excitation at 405 nm.

The InP from additional rounds of SILAR deposition enhance the concentration-normalized absorbance of the particles (**Figure 4a, Figure S8**), as is appropriate given the larger volume of InP per particle. The distinct 1S peaks seen in the photoluminescence excitation (PLE) spectra of the InP QDs emitting in the visible wavelength regime (**Figure 4b**) are indicative of quantum confinement. The QD samples exhibited absolute quantum yields up to 38%, as measured in an integrating sphere (Horiba Quanta-ϕ) (**Table S2**). For the three reddest emitters, which exhibited significant PL outside the calibrated range of our integrating sphere, we used the directly measured absolute QY and full PL spectrum to extrapolate the QY of the redder region using the relative area under the curve.

The weighted average photoluminescence lifetimes of the InP QDs, determined using a tri-exponential fit, are between 80 and 140 ns, similar to previously observed lifetimes seen for InP QDs (**Figure 4c, Table 1, Table S3**).^20, 22, 23, 46, 47^ The inverted InP QD emission peaks are broad compared to InP core-based systems, however, and the largest particles exhibit red-tail emission > 1000 nm, which is beyond the bulk bandgap of InP. Investigation of the fluorescent lifetimes of different spectral regions of the QD555 and QD675 emission peaks (*i.e*., blue tail, peak, and red tail emission) reveal wavelength region-dependent differences in the fluorescent lifetimes, as shown in **Figure S9**. Specifically, the weighted average PL lifetimes increase with increasing emission wavelength (*i.e*., 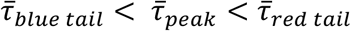). This trend is particularly evident for the transition from the emission spectra peak region to the red tail, where the average lifetimes increase from 102 ns to 223 ns and 147 ns to 426 ns for QD555 and QD675, respectively **(Table S4)**. The breadth of the emission peak and differences in the lifetimes suggest that there may be different emission mechanisms between these spectral regions of the inverted QD system or, alternatively, a heterogeneous particle population. While there is precedent for QD systems with more than one emission mechanism,^48, 49^ further studies will be required to ascertain the precise source of the wavelength dependent lifetimes in this QD system.

Notably, the NIR-emitting particles exhibit significant absorption and PLE > 600 nm (**Figure 4**). Efficient excitation with red and NIR light ensures that penetrating wavelengths can be used for *in vivo* photoluminescence, enhancing the usefulness for preclinical imaging. Through this inverted InP QD synthesis, we have achieved extended emission tunability compared to what has been previously published for InP core-based QDs, yielding multiple emitters in the first optical tissue window. In order to generate bright NIR emitters for *in vivo* imaging of sentinel lymph nodes, QD675, QD770, and QD845 were Zn-TOP treated to enhance their brightness. As a result of this treatment, the QY of the QDs increased, but the emission peaks of the two longer wavelength emitters blue shifted to QD720 and QD790, respectively (**Table S2)**. Following a previously published protocol, QDs were rendered water soluble through encapsulation in lipid-PEG micelles using 1,2-distearoyl-sn-glycero-3-phosphoethanolamine-N-[methoxy(polyethylene glycol)] (ammonium salt) with 5000 Da PEG (DSPE-PEG5k, Avanti Polar Lipids, Inc.).^50, 51^ The water soluble samples largely retained their pre-coating brightness, exhibiting QYs between 11 and 18%, consistent with similarly treated commercial QD705s (Thermo Fisher) (**Table S2)**.

To demonstrate multiplexed *in vivo* imaging with the NIR-emitting InP QDs, the three Zn-TOP-treated samples were encapsulated in in DSPE-PEG5K, buffer exchanged into phosphate buffered saline (PBS), and sterile filtered. QDs were subcutaneously injected into a BALB/cJ mouse in three distinct sites to visualize lymphatic drainage. QD675 and QD790 were injected into the right and left hock, respectively. QD720 was injected into the right second mammary fat pad (**Figure 5a**). The mouse was imaged using an In Vivo Imaging System (IVIS Spectrum; Perkin Elmer) at several time points post injection (p.i.), and the resulting images were analyzed and unmixed using the Living Image software (Perkin Elmer) to show the emission from each QD. At 28 minutes p.i., drainage from the initial site of injection to local lymph nodes (LNs) was observed for all three of the QDs (**Figure 5b)**. One-hour p.i., lymphatic drainage is visible in the collecting lymphatic vessel extending from the left inguinal LN to the left axillary LN (**Figure 5c**). We included an inset at higher thresholding that shows the inguinal LN itself, since the low thresholding that enables visualization of the lymphatic drainage vessel causes saturation of the region **(Figure 5, Figure S10)**. Visualization of QD720 and QD790 in the lymph nodes was possible at up to 52 hours p.i. when the animal was sacrificed and the body cavity exposed to confirm the co-localization of QDs with LNs (**Figure S11**). QD720 and QD790 were located in the axillary LNs on their respective sides of injection. QD790 was also seen in several LNs near the inguinal node as well as the inguinal LN itself. QD675 was not detected during the post-mortem analysis.

**Figure 5.**
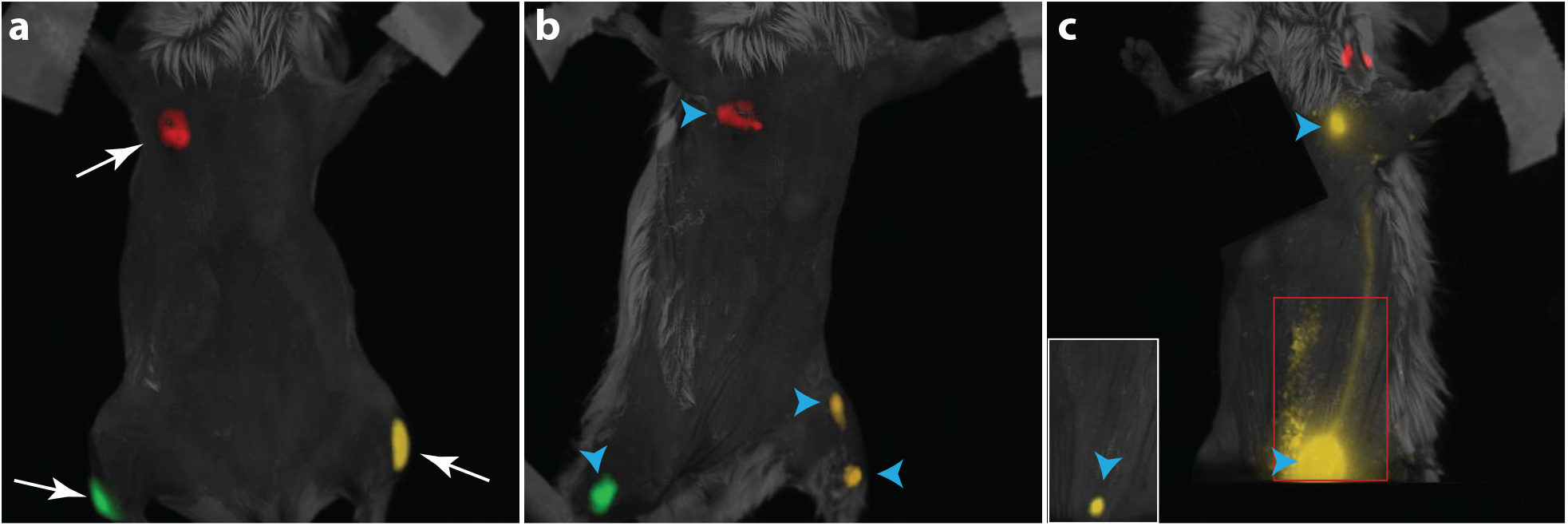
Spectrally unmixed images of a BALB/cJ mouse injected with three distinct QD solutions: QD675 (green; left hock injection), QD790 (gold; right hock injection), and QD720 (red, left armpit). (**a**) Injection sites imaged immediately post injection (p.i.). (**b**) Localization of QD675 in the left inguinal lymph node (LN), QD720 in the axillary LN, and QD790 in the right inguinal LN and popliteal LN 48 min p.i. (**c**) Lateral image 1 hr p.i. shows QD790 lymphatic drainage from the inguinal LN to the axillary LN; inset of the boxed region with higher thresholding shows the contrast localized in the inguinal LN without image saturation. Images taken on Perkin Elmer IVIS Spectrum and analyzed using Living Image software. White arrows indicate injection sites, while blue arrowheads indicate LNs.

In summary, through multi-component InP SILAR deposition in the presence of ZnSe cores, we synthesized indium phosphide-based QDs that exceed the previous NIR emission limits for this important cadmium-free composition. These small, bright nanocrystals comprising only In, P, Zn, S, and Se can be tuned to emit throughout the first optical tissue window, presenting an opportunity for multiplexed fluorescent imaging in the NIR, with QYs consistent with commercially available cadmium-based QDs. Subsequent work will optimize the synthesis to narrow the emission bandwidths, further improve brightness, and tailor the colloidal coating for targeted imaging.

## ASSOCIATED CONTENT

### Supporting Information

A full description of the nanoparticle synthesis and coating procedure as well as additional characterization results are included in the Supporting Information. The Supporting Information is available free of charge on the ACS Publications website.

## Author Contributions

A.M.S and A.M.D. conceived of the project. A.M.S., A.Y.N., R.T., J.C.K., M.C and A.M.D. designed experiments. A.M.S. performed the synthesis and micelle encapsulation of QDs, as well as MP-AES experiments. A.Y.N. and A.M.S. carried out STEM and EDS experiments. A.M.S., J.C.K. and R.T. performed optical analysis. J.P.C and K.H. performed STEM sizing. A.M.S. and D.J. performed the lymph node imaging experiments. A.P performed EMM calculations. A.M.S. and A.M.D. wrote the manuscript with comments and editing from co-authors, and all authors have given approval to the final version of the manuscript.

## ACKNOWLEDGMENT

The research reported in this publication was supported by the National Institute of General Medical Sciences of the National Institutes of Health under award number R01GM129437. This work was supported in part through a Boston University Clinical and Translational Science Institute (CTSI) KL2 Fellowship to A.M.D. (1KL2TR001411). Financial support for A.M.S. provided in part by a BUnano Cross-Disciplinary Fellowship. Financial support for J.C.K. provided by Training Grant NIH/NIGMS T32 GM008764 and by the National Science Foundation Graduate Research Fellowship (NSF-GRFP) under Grant No. DGE-1840990. Financial support for M.C. provided through the Clare Boothe Luce (CBL) Program from the Henry Luce Foundation. This work was performed in part at the Harvard University Center for Nanoscale Systems (CNS), a member of the National Nanotechnology Infrastructure Network (NNIN), which is supported by the National Science Foundation under NSF award no. ECS-0335765. CNS is part of Harvard University. IVIS imaging performed at the BU Metabolic Phenotyping and Imaging Core, supported in part by grant 1S10RR024523-01. Core facilities provided by the BU Photonics Center and Division of Materials Science and Engineering include the CARY5000, Technai Osiris (S)TEM/EDS, and Bruker D2 Phaser instruments. A.P. acknowledges the support by the Los Alamos Laboratory Directed Research and Development (LDRD) funds.

## Table of Contents

The photoluminescence of InP-based quantum dots (QDs) is extended into the first optical tissue window by adopting an inverted structure. The ZnSe/InP/ZnS core/shell/shell particles exhibit InP shell thickness-dependent tunable emission throughout visible and near infrared (NIR) wavelength ranges. These toxic metal-free nanoparticles both absorb and emit in the NIR making them a promising contrast agent for bioimaging.

For TOC only:

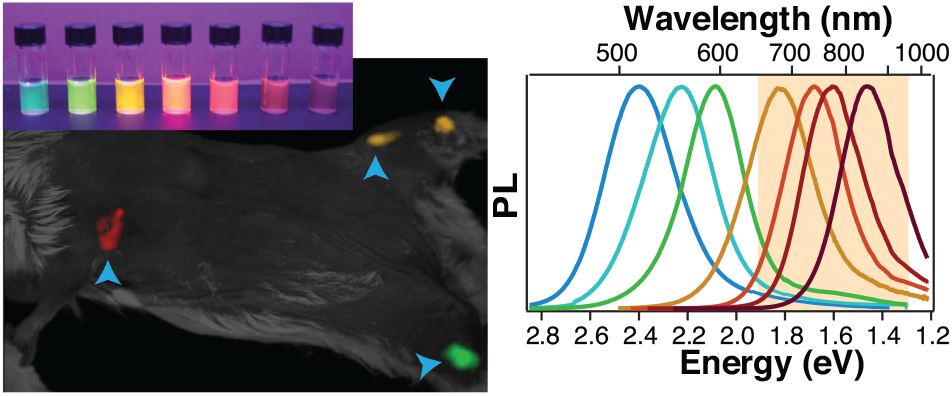

