## Supplementary Information for "Extending the Near Infrared Emission Range of Indium Phosphide Quantum Dots for Multiplexed *In Vivo* Imaging"


### 1 Experimental Section

#### 1.1 Materials

Zinc acetate (99.99%), oleic acid (technical grade, 90%), selenium pellets <5 mm (trace metals grade, ≥ 99.99%), 1-octadecene (ODE, technical grade, 90%), oleylamine (technical grade, 70%), diphenylphosphine (DPP, 98%), toluene (anhydrous, 99.8%), hexane (for HPLC, ≥ 98.5%), Trioctylphosphine (TOP, 97%), and indium (III) acetate ($In(Ac)_3$, 99.99%) were purchased from Sigma-Aldrich. Tris(trimethylsilyl)phosphine ($TMS_3P$, 98%) and ethanol (anhydrous, histological) were purchased from Fisher Scientific, and sulfur (refined, 99.5%) was purchased from ACROS Organics. 1,2-distearoyl-sn-glycero-3-phosphoethanolamine-N-[methoxy(polyethylene glycol)-5000] (ammonium salt) (DSPE-PEG5k) was purchased from Avanti Polar Lipids, Inc. Quartz glass cuvettes were purchased from Starna Cells Inc. All air-sensitive materials were stored and handled in a glovebox under argon.

#### 1.2 Precursor Synthesis

##### *1.2.1 Zinc Oleate*

The 0.2 M zinc oleate precursor was synthesized by reacting zinc acetate and oleic acid in ODE. A 250 mL, 3-neck, round-bottom flask containing 6.59 g zinc acetate (35.9 mmol), 42 mL oleic acid (120 mmol), and 108 mL ODE was mounted on a Schlenk line, connected to a thermocouple, and sealed with a septum. The flask was placed under vacuum, heated to 120 °C, and stirred continuously. Vigorous boiling was observed, due to the release of acetic acid during the conversion of zinc acetate to zinc oleate. Once boiling ceased, the reaction flask was backfilled with argon and the reaction temperature was increased to 180 °C for 30 minutes under dynamic argon flow to ensure that the conversion reaction had gone to completion. The temperature was then reduced to 120 °C and the solution placed under vacuum for 1 hour to remove any residual acetic acid. The final zinc oleate precursor was stored under argon at room temperature, forming a waxy solid. When needed, the zinc oleate was heated at 80 °C under active argon flow to liquefy the precursor and allow for withdrawal with a needle and syringe.

#### *1.2.2 Zinc-TOP solution*

The 0.2 M zinc TOP solution was synthesized as described for zinc oleate in section 1.2.1, with the modification of substituting TOP as a reaction solvent in the place of ODE. Unlike 0.2 M zinc oleate, the 0.2 M Zinc-TOP solution is not solid at room temperature and therefore was not heated prior to use.

#### *1.2.3 Diphenylphosphine Selenide*

Diphenylphosphine selenide (DPP:Se) was synthesized as previously described. [1] In an argon-filled glovebox, 1.58 g of selenium pellets (20.0 mmol) were added to a 3-neck reaction flask containing a magnetic stir bar. Next, 3.48 mL DPP (19.6 mmol) and 25 mL anhydrous toluene were added to the reaction flask. A condenser and stopcock were added to the 3-neck reaction flask in addition to a thermocouple; finally, the 3-neck reaction flask was sealed with a septum to maintain an inert atmosphere. The reaction flask was brought out of the glovebox, mounted on a Schlenk line, and placed under active argon flow. The reaction was heated until it started gently boiling, and was refluxed at this temperature (114 °C) with a Findenser Super Air Condenser for 16 hours. Next, the reaction flask was transferred to a rotary evaporator and the toluene gently evaporated, yielding large flakey crystals of DPP:Se. These crystals were collected and washed with cold toluene over a Buchner funnel using filter paper. The resulting purified DPP:Se was a clear white powder that was stored in the glovebox under an inert atmosphere.

#### *1.2.4 Indium Oleate*

The 0.2 M indium oleate precursor was synthesized by reacting oleic acid with indium acetate in ODE. In an argon-filled glovebox, 5.84 g indium acetate (20.0 mmol) were added to a 250 mL, 3-neck, round-bottom flask. The flask was removed from the glovebox and mounted on a Schlenk line. Next 23.2 mL of Oleic acid (66.1 mmol) and 76.9 mL of ODE were added to the reaction flask before it was connected to a thermocouple and sealed using a septum. The flask was placed under vacuum while the solution was stirred and heated to 120 °C. The solution exhibited vigorous bubbling during the conversion of indium acetate to indium oleate as acetic acid was released. Once the solution ceased to boil, the reaction flask was placed under active argon flow and the reaction temperature was increased to 160 °C for 20 min to ensure that the conversion reaction had gone to completion. The temperature was returned to 120 °C, and the reaction was placed under vacuum for 1 hour to remove any residual acetic acid. The indium oleate was stored at room temperature under argon. When needed, the indium oleate precursor solution was heated to 40 °C under active argon flow to reduce its viscosity for improved dispensing with a needle and syringe.

#### *1.2.5 Sulfur Solution*

A 0.2 M sulfur solution in ODE (S8/ODE) was synthesized by adding 0.64 g of sulfur (20.0 mmol) and 100 mL ODE to a 250 mL, 3-neck, round-bottom flask containing a magnetic stir bar while in a glove box under argon. A thermocouple was added to the flask, which was then removed from the glovebox and mounted on a Schlenk line. Next the reaction solution was heated to 120 °C and placed under vacuum for 2 hours until the sulfur fully dissolved. The reaction flask was backfilled with argon and the temperature was then reduced to 80 °C and allowed to stir overnight. The 0.2 M S8/ODE solution was kept at 80 °C to prevent precipitation of sulfur from the solution.

#### *1.2.6 Selenium Solution*

A 0.2 M stock solution of TOP:Se was prepared by adding 1.58 g of selenium pellets (20.0 mmol) and 100 mL TOP (224 mmol) to a 250 mL 3-neck round-bottom flask containing a magnetic stir bar while in an argon glovebox. A condenser and temperature controller were added to the flask and it was sealed using a septum. The flask was then removed from the glovebox and mounted on a Schlenk line, where it was vacuumed for one hour and backfilled with argon. The solution was heated to 80 °C and stirred and kept at that temperature overnight. After the complete dissolution of the selenium pellets, the flask was removed from heat.

### 1.3 Nanoparticle Synthesis

#### *1.3.1 ZnSe cores*

Zinc selenide (ZnSe) cores were synthesized according to a protocol previously described in the literature. [2] Briefly, 6 mL of 0.2 M zinc oleate, was added to a 100 mL, 3-neck, round-bottom flask under active argon flow. The solution was heated to 80 °C, and 0.2 mL DPP was added to increase the reactivity of the zinc oleate. DPP was allowed to react with zinc oleate for 10 minutes at 80 °C. Next, 4 mL of 0.15 M DPP:Se dissolved in toluene was added to the reaction flask. Vacuum was applied for 5 min to remove the toluene before the solution was placed under active argon flow. The temperature was increased at a rate of 10 °C per min until a temperature of 220 °C was reached, at which point the reaction was allowed to proceed for 15 min to completion. The resulting cores were brought into a glovebox and centrifuged to remove excess zinc. After centrifugation, the supernatant was transferred to a 50 mL conical tube, and the pellet discarded. The supernatant was precipitated by adding a two-fold excess of anhydrous hexanes to the ZnSe particles in ODE followed by a four-fold excess of anhydrous ethanol, at which point the supernatant became cloudy. The cloudy solution was centrifuged for 10 minutes to form a solid pellet of ZnSe cores. The supernatant containing unreacted precursors was discarded and the pelleted cores redispersed in ODE and stored under argon in the glovebox until needed.

ZnSe cores were characterized using transmission electron microscopy (TEM) and absorbance spectroscopy (both described below). The core diameter and molar extinction coefficient were determined using the 1S peak position in conjunction with empirical fit equations. [3] The calculated molar extinction coefficient and diameter of the core were $\varepsilon_{355} = 1.24\times10^5$ $M^{-1}$ $cm^{-1}$ and d= 2.06 nm respectively.

#### *1.3.2 ZnSe/InP/ZnS core/shell/shell*

Indium phosphide (InP) shells were deposited on the ZnSe cores using a modified successive ion layer adsorption and reaction (SILAR) scheme. [4] Six of the seven QDs presented in the manuscript (QD515, QD555, QD595, QD675, QD740, and QD770) were synthesized in the same manner on the same ZnSe cores, but with varying numbers of rounds of SILAR, resulting in distinct thicknesses of the InP shell layer and photoluminescence emission wavelengths. Each of these samples was synthesized in its own reaction flask, starting with 9 mL ODE and 1 mL oleylamine, stirred under vacuum at 120 °C for 1 hour to remove air and residual water. Each reaction flask was put under active argon flow, cooled to room temperature, and 150 nmol of 2.06 nm diameter ZnSe cores were added via syringe. The solution was heated to 150 °C, after which a volume of 0.2 M $In(OA)_3$ sufficient to add one monolayer of indium to the ZnSe core was added and allowed to react for 15 minutes. The same volume of 0.2 M tris(trimethylsilyl)phosphine $(TMS)_3P$ was added dropwise, and the reaction was heated to 178 °C for a 1 hour anneal. The volume of 0.2 M $In(OA)_3$ and $(TMS)_3P$ needed to add the second atomic monolayer (and subsequent layers) was similarly calculated, taking the expected increase in particle size into account. The required volume of 0.2 M $In(OA)_3$ was added dropwise to the solution followed by a 1 hour anneal at 178 °C; $(TMS)_3P$ was subsequently added, followed by a 1 hour anneal. The process of cation addition and anneal followed by anion addition and anneal was repeated to generate a series of InP shell thicknesses (2, 3, 4, 7, 10, and 13 SILAR iterations for QD515, QD555, QD595, QD675, QD740, and QD770, respectively). To form a zinc sulfide (ZnS) cap on our ZnSe/InP QDs, 0.2 M zinc oleate was added to the reaction solution dropwise at 178°C. After addition, the reaction was heated to 220°C followed by a 1 hour anneal time. Then sulfur dissolved in ODE (S8/ODE, 0.2 M), synthesized as described in section 1.2.4, was likewise added dropwise and allowed to anneal for 1 hour. Two or three rounds of zinc and sulfur deposition were used for these samples.

The final sample presented here, QD845, was similarly synthesized with multiple minor modifications. Specifically, a different ZnSe core was used (3.1 nm diameter), the initial reaction solvent comprised 2 mL oleylamine and 8 mL ODE, cation anneals were 2.5 hours long, the anion precursor was more dilute at 0.1 M $(TMS)_3P$. Sixteen iterations of SILAR additions were performed, and significant samples were taken at thinner shells for reaction tracking, but SILAR addition volumes were adjusted to take this particle removal into account.

#### *1.3.3 Zinc TOP-treatment*

Zinc-TOP treatment of QDs was carried out by first precipitating QDs from the reaction solution by adding a mixture of hexanes and ethanol (1:3) followed by centrifugation to generate a nanocrystal pellet. Argon was then used to fill the centrifugal tube containing the NC pellet with an inert atmosphere and 1.5 mL of Zn-TOP solution was added to the tube after which argon was again flowed over the tube before closing the tube. The tube was then vortexed and placed in a heated sonication bath (45°C) for 15 minutes after which it was again vortexed and placed back in the sonication bath. This process was repeated until the NC pellet had fully dispersed in the Zinc-TOP solution, normally 2-3 iterations, after which the Zinc-TOP solution containing the QD were transferred to 2-dram borosilicate glass vials, containing a further 1.5 mL of Zinc-TOP solution, after which the vials were once again flushed with argon and sealed using polytetrafluoroethylene (PTFE) lines caps. These vials were vortexed for 30s to allow for adequate mixing and then placed on a temperature regulated hot plate, where the temperature was ramped from room temperature to 240°C. The QD solutions were incubated in the Zinc-TOP solutions for 3 hours at 240°C, after which the solutions were removed from the hot plate and cooled to room temperature. The solutions were then precipitated in a mixture of hexanes and ethanol (1:4) and suspended in the desired solvent.

### 1.4 Micelle encapsulation

QDs were encapsulated according to literature, with modifications. In a tared tube, QDs were precipitated with hexane and ethanol (1:3) followed by centrifugation to generate a nanocrystal pellet. [5] The supernatant was discarded, and the QD sample was allowed to dry for 5 minutes to remove any remaining solvents. The tube was weighed to calculate the mass of QDs precipitated. The pellet was redispersed in chloroform at a concentration of 4 mg/mL. DSPE-PEG2k was dissolved in chloroform (15 mg/mL) and added to the QD sample solution at a QD:DSPE-PEG2k ratio of 1:4.5 (by weight), and allowed to interact for 10 min. This solution was added to a pear-shaped flask and rotary evaporated at 70 °C until a thin film formed. Heated deionized (DI) water (15 mL) and marbles at 70°C were rapidly added to the thin film followed by vigorous swirling of the flask to form micelles. The sample was recovered using a syringe and was passed through a 0.22 µm syringe filter to remove any aggregates. The resulting QD micelles were concentrated using centrifugal molecular weight cut off filters (100 kDa).

### 1.5 Bandgap calculations using core/shell structures effective mass model (EMM)

The electronic structures of both the traditional and inverted heterostructures were calculated using a two-band EMA applied to the core/single-shell heterostructures (i.e., InP/ZnSe and ZnSe/InP). The effect of the high bandgap ZnS cap (second shell) is negligible and was not accounted for in the calculations. The confinement potential is of Type I in character and has been parametrized as follows. InP bandgap energy is set to the bulk value $E_g$ = 1.34 eV. At the interface, conduction (valence) band energy step is assigned to $U_e$ = 0.3 eV ($U_h$ = -1.08 eV) defining the electron (hole) delocalization energy. The 1S exciton energy is evaluated as a sum of the electron and hole 1S state energies due to the confinement potential (kinetic energies) plus their Coulomb binding energy (perturbative corrections). For InP and ZnSe, electron (hole) effective masses are set to $m^*_e = 0.073m_0$ ($m^*_h = 0.64m_0$) and $m^*_e = 0.15m_0$ ($m^*_h = 0.53m_0$), respectively. [6,7] Values of high frequency dielectric constants for InP and ZnSe are set to 9.61 and 5.6, respectively. [7]

### 1.6 X-ray diffraction (XRD)

ZnSe cores and ZnSe/InP/ZnS QDs were precipitated from ODE by mixing the nanoparticle laden ODE solution with a two-fold excess of hexane followed by twice as much ethanol. The precipitating nanoparticles were pelleted through centrifugation and the supernatant discarded. The QDs were resuspended in hexanes and drop-cast onto a zero-background silicon sample holder. A Bruker D2 Phaser system was used to acquire powder x-ray diffraction data using a coupled theta-2 theta scan. Reference diffraction lines for ZnSe, InP, and ZnS were generated using the free software PowderCell2.4 downloaded from the Federal Institute for Materials Research and Testing (Bundesanstalt für Materialforschung und –prüfung, BAM, Berlin, Germany). These reference lines are representative of the idealized positions and intensities of the crystallographic peaks for each material's bulk diffraction.

### 1.7 Optical spectroscopy

#### *1.7.1 Photoluminescence emission spectroscopy*

Photoluminescence (PL) spectra were recorded with a Nanolog® Spectrofluorometer with a Synapse CCD camera and iHR320 fully automated spectrometer (HORIBA Jobin Yvon). QDs suspended in hexanes were loaded into quartz cuvettes and inserted into the sample chamber. Samples were excited at 375 nm, and the resulting emission spectra was collected with a 0.5 second integration time and corrected for lamp power and detector sensitivity using the supplied correction files.

#### *1.7.2 Photoluminescence excitation spectroscopy*

Three-dimensional scans of excitation wavelength and emission intensity were measured on the Nanolog® Spectrofluorometer with a Synapse charge coupled device (CCD) camera and iHR320 fully automated spectrometer (HORIBA Jobin Yvon). QDs suspended in hexanes were loaded into quartz cuvettes and inserted into the sample chamber. The integration time was set to 5 seconds, and spectra were corrected for lamp power and detector sensitivity using the supplied correction files. Full emission spectra were collected per each excitation wavelength, testing from 350 nm excitation to 902 nm in 4 nm steps.

#### *1.7.3 Photoluminescence lifetime spectroscopy*

Photoluminescence Lifetime measurements were acquired through Time-Correlated Single Photon Counting (TCSPC) on a LifeSpec II spectrophotometer. QDs suspended in hexanes were loaded into quartz cuvettes and inserted into the sample chamber. Samples were excited with a 405 nm EPL picosecond pulsed diode laser using a 1 microsecond pulse period. For each sample, the measurement continued until 20,000 photons were collected in the highest-intensity bin to ensure accurate fits. Photons were collected at the emission peak with a bandwidth window of 15-50 nm, depending on the full width at half maximum (FWHM) of the sample. To assess lifetimes in different regions of the emission spectra, photons were collected only from the blue tail, red tail and peak regions with consistent 25 nm bandwidth windows placed within each of these regions. After collection, data was analyzed with F980 software (Edinburgh Instruments) by fitting the tail to the tri-exponential equation $R(t) = A_1 e^{-t/\tau_1} + A_2 e^{-t/\tau_2} + A_3 e^{-t/\tau_3}$ , where $A_n$ is the percent weight of each exponential. The average photoluminescence lifetime, $\bar{\tau}$, was calculated using equation 1. [8, 9]

$$\bar{\tau} = \sum_n \frac{A_n \tau_n^2}{\sum_m A_m \tau_m} \qquad (1)$$

#### *1.7.4 Absolute quantum yield determination*

Absolute quantum yield values were obtained using the spectrofluorometer described above in conjunction with a Quanta-phi six-inch integrating sphere (HORIBA Jobin Yvon). Cuvettes were loaded with hexanes in order to generate a blank reference following excitation at 400 nm with an integration time of 0.3 seconds. QD samples were diluted directly into the blanked cuvettes and measured with identical conditions. The absolute quantum yield (*i.e.*, the photons emitted divided by the photons absorbed) was determined using the Horiba software.

#### *1.7.5 Absorbance spectroscopy*

Absorption spectra were recorded using a CARY5000 UV/Vis Spectrophotometer from Agilent, Inc. Controlled volumes of the reaction solutions were diluted in hexanes and the absorbance of this "raw" sample measured. These controlled dilutions were used to determine the concentration-normalized absorbance of the discrete QD samples, similarly to a previously published report. [10] We estimated the concentration of the QDs in the raw solution by assuming that the final QD concentration is equal to the initial concentration of ZnSe cores adjusted for precursor addition volumes.

To ensure that the increase in absorbance does not stem from unreacted precursor, the absorbance of cleaned samples (*i.e.*, QDs precipitated and resuspended in hexanes) was also measured and compared to the spectra of the raw samples. Washed QDs in hexanes were measured with a wavelength step of 1 nm and an integration time of 0.1 second. In order to normalize for the particle concentration of the cleaned sample (*i.e.*, to account for a loss of QDs during the purification), we concentration-normalized the "clean" spectra by scaling it to the absorbance spectra of the raw samples.

### 1.8 Microwave plasma atomic emission spectroscopy (MP-AES) for elemental analysis

Samples were prepared for MP-AES elemental analysis by repeated precipitation by the addition of hexanes and ethanol (1:3) followed by centrifugation, a brief drying step where the centrifugal tubes were placed top down on a kimwipe to allow the solvents to escape the tube, and then resuspension in hexanes, followed precipitation in ethanol. This process was repeated five times to ensure that no residual precursor solution was present, as this could confound analysis. After the fifth precipitation the samples were suspended in chloroform and transferred to trace metal clean 2-dram vials and allowed to evaporate under mild heating, 40°C, overnight on a hot plate. To the 2-dram vials containing the dry QD films was added 300 µL of trace metal grade aqua regia, the vials were immediately capped with PTFE lined caps (PTFE caps are a requirement as other caps rapidly disintegrate under QD digestion conditions) and transferred to a hot plate that was already set to 80°C and allowed to digest at 80°C for 24 hours. The resulting solutions were optically transparent, with some small optically clear oily inclusions that were likely the surface ligands of the QDs and ranged from clear to tinged slightly yellow in color. Ultrapure water was then added to these samples in sufficient amounts to bring the total acid concentration in the solution to 7% to prepare the samples for analysis.

To ensure the instrument torch was equilibrated prior to analysis, the MP-AES (Agilent 4210) was warmed up for one hour prior to start of analysis. A standard curve was generated using solutions containing known concentrations of elements, using high quality elemental standards (High Purity Standards). The samples were run using the manual configuration of the MP-AES, with four instrument measurement replicates collected per element, per sample.

### 1.9 Transmission electron microscopy

Transmission electron microscopy (TEM) images were obtained using a JEOL 2100 system operating at 200 keV. Samples were prepared by drop-casting QDs suspended in hexanes onto a standard copper TEM grid. The TEM grid was subsequently cleaned by successive washing with hexanes, ethanol, and DI water, followed by gentle heating to evaporate residual water.

Determination of the particle sizes from TEM images was performed manually using ImageJ software.

Scanning transmission electron microscopy (STEM) imaging and energy dispersive spectrometry (EDS) mapping were performed on a Tecnai Osiris TEM from FEI, Inc. operated in STEM mode equipped with a windowless four-quadrant Super X EDS detector for analytical work. Samples for STEM/EDS experiments were prepared on ultra-thin carbon copper grids following the steps used for standard TEM imaging described above. Prior to examination in TEM, the samples were plasma cleaned for 15 sec in a 1040 plasma cleaner from Fischione Instruments, Inc. using a shielded holder accessory for plasma cleaning sensitive samples. To minimize potential beam damage to indium atoms in the inner shell of the QD structures, both imaging and elemental mapping were always performed on fresh locations with X-ray count collection times not exceeding about 300 sec, when the beam damage to the examined area became noticeable.

#### 1.10 Wide-field mouse imaging

TOP treated and micelle encapsulated QDs were buffer exchanged three times with 1X phosphate buffered saline, and sterile filtered using a 0.1 µm filter. 50 µL of sterile solutions of QD790, QD720, and QD675 were loaded into insulin injection syringes (BD-Safety glide insulin syringe and needle, 29G). QD solutions were injected into the left leg (QD790-hock injection), right axilla (QD720), and right leg (QD675-hock) respectively, in that sequence, while the anesthetized and chemically shaved mouse was on the imaging platform. Optical imaging using the IVIS Spectrum imaging system (Perkin Elmer) commenced immediately and was repeated frequently while altering the mouse position and optimizing the imaging settings. The mouse position and image settings (exposure time) were optimized to reveal different biological structures, including lymphatic vessels and primary and secondary lymph nodes draining the site of injection. Images were gathered during three separate imaging sessions: the first session encompassed time points immediately post injection and up to two hours p.i., a second session spanning 32-33 hours p.i., and a final session spanning 54-55 hours p.i. ending in the imaging of the dissected animal to show the colocalization of the signal with the lymph node.

All images were analyzed and unmixed using the Living Image software (Perkin Elmer). The number of components included for unmixing was determined using the built-in principal component analysis suite in Living Image.

### 2 Supplemental Data

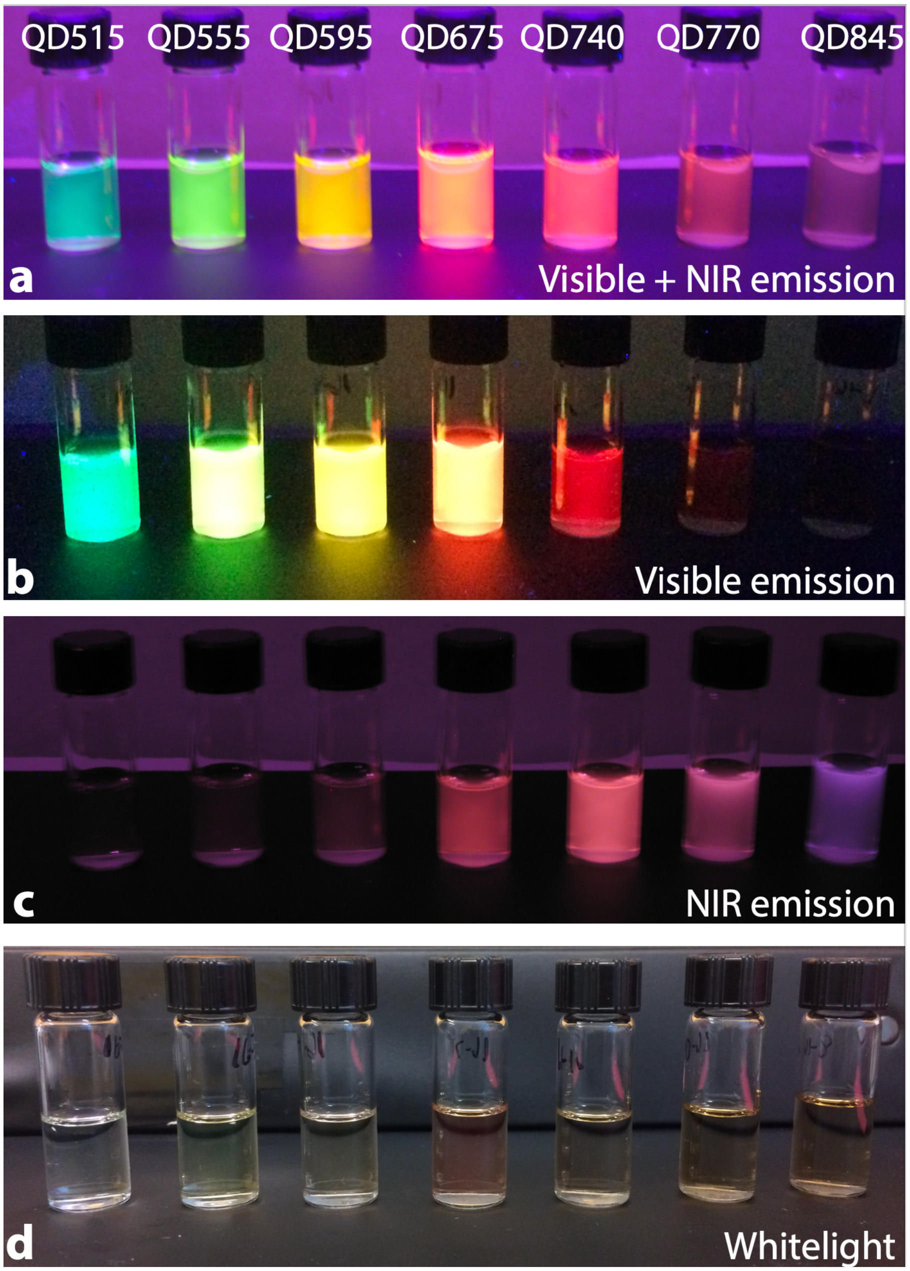


**Figure S1. Photos of QDs**. a) QDs under UV illumination; digital camera internal IR-blocking filter removed to show visible and NIR emission. b) QDs under UV illumination, with standard IR-blocking filter. c) QDs under UV illumination, internal IR-blocking filter removed, and 720nm long pass filter used to detect NIR emission. d) QDs under white light illumination, with standard IR-blocking filter

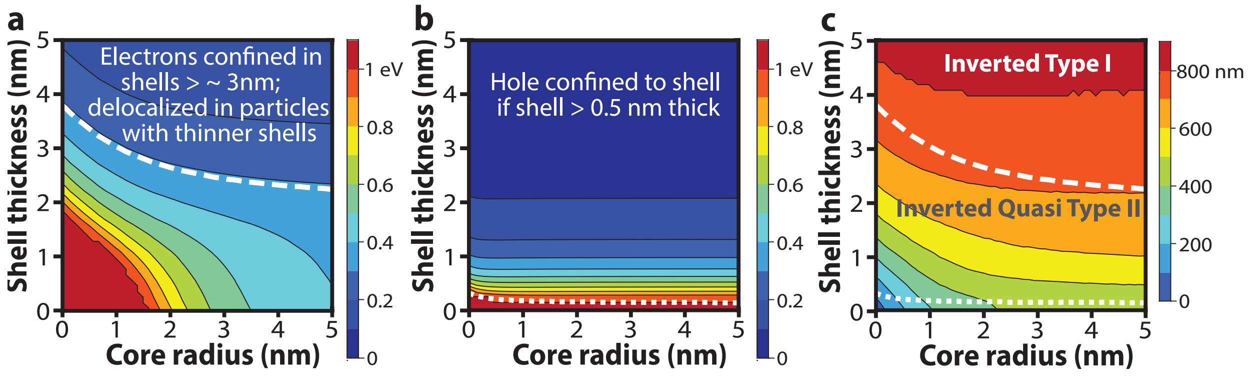


**Figure S2. Two-band effective mass model calculations of ZnSe/InP heterostructure** incorporating perturbative corrections for Coulomb binding energy. (**a**) Electron 1S state energies as a function of core radius and shell thickness in eV. (**b**) Hole 1S state energy as a function of core radius ad shell thickness. (**c**) Predicted 1S bandgap as a function of core radius and shell thickness in nm. In all plots, the dashed white line indicates the transition from delocalized to confined electron wave function (defined as 90% localized); the dotted white line indicates the transition from delocalized to confined hole wave function (defined as 90% localized).

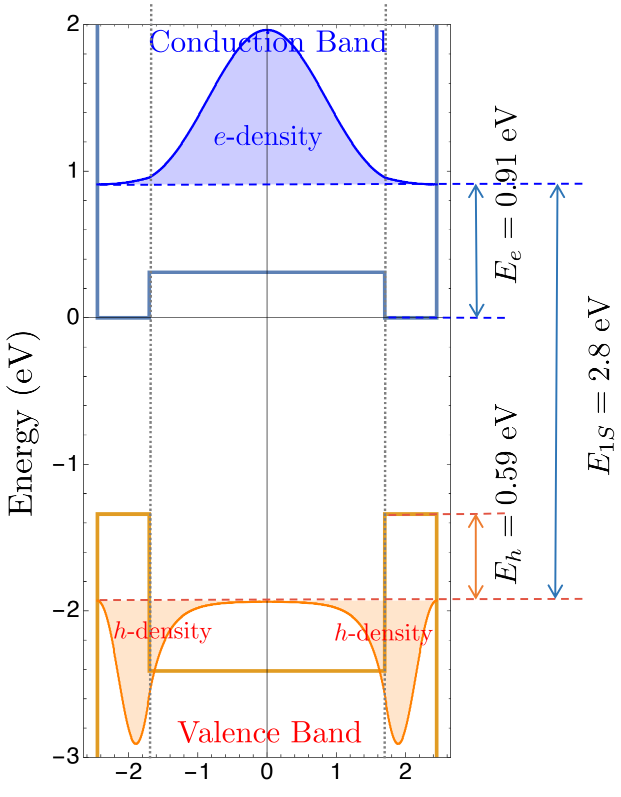


**Figure S3.** Electron and hole density plot for ZnSe/InP core/shell with a 1.7 nm radius (3.4 nm diameter) core and 0.75 nm thick shell.

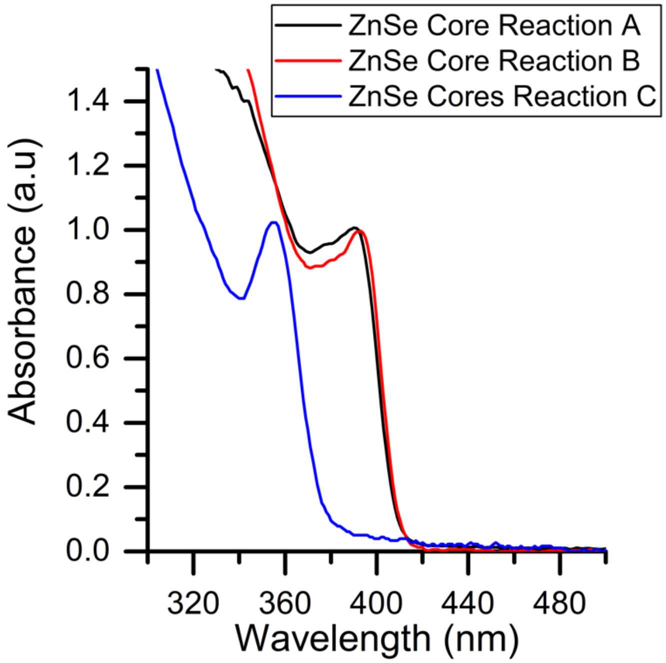


**Figure S4. Absorbance spectra of ZnSe cores**. ZnSe core samples exhibited 1S peaks at (A) 393 nm (black line), (B) 390 nm (red line), and (C) 356 nm (blue line), corresponding to core diameters of 3.1, 3.1, and 2.1 nm, respectively.

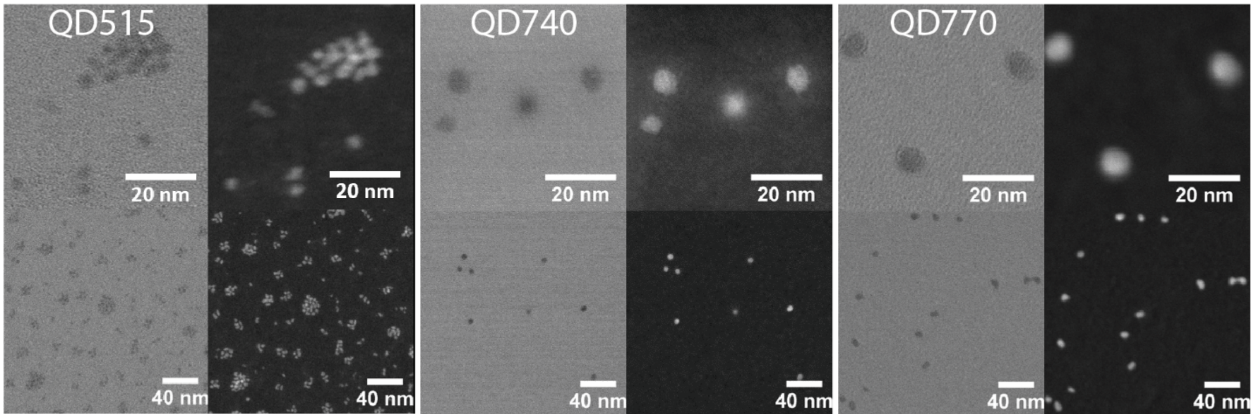


**Figure S5. Bright-field (BF; left) and high-angle annular dark-field (HAADF; right) STEM images** of QD515, QD740, and QD770 showing particle relative size and morphology.

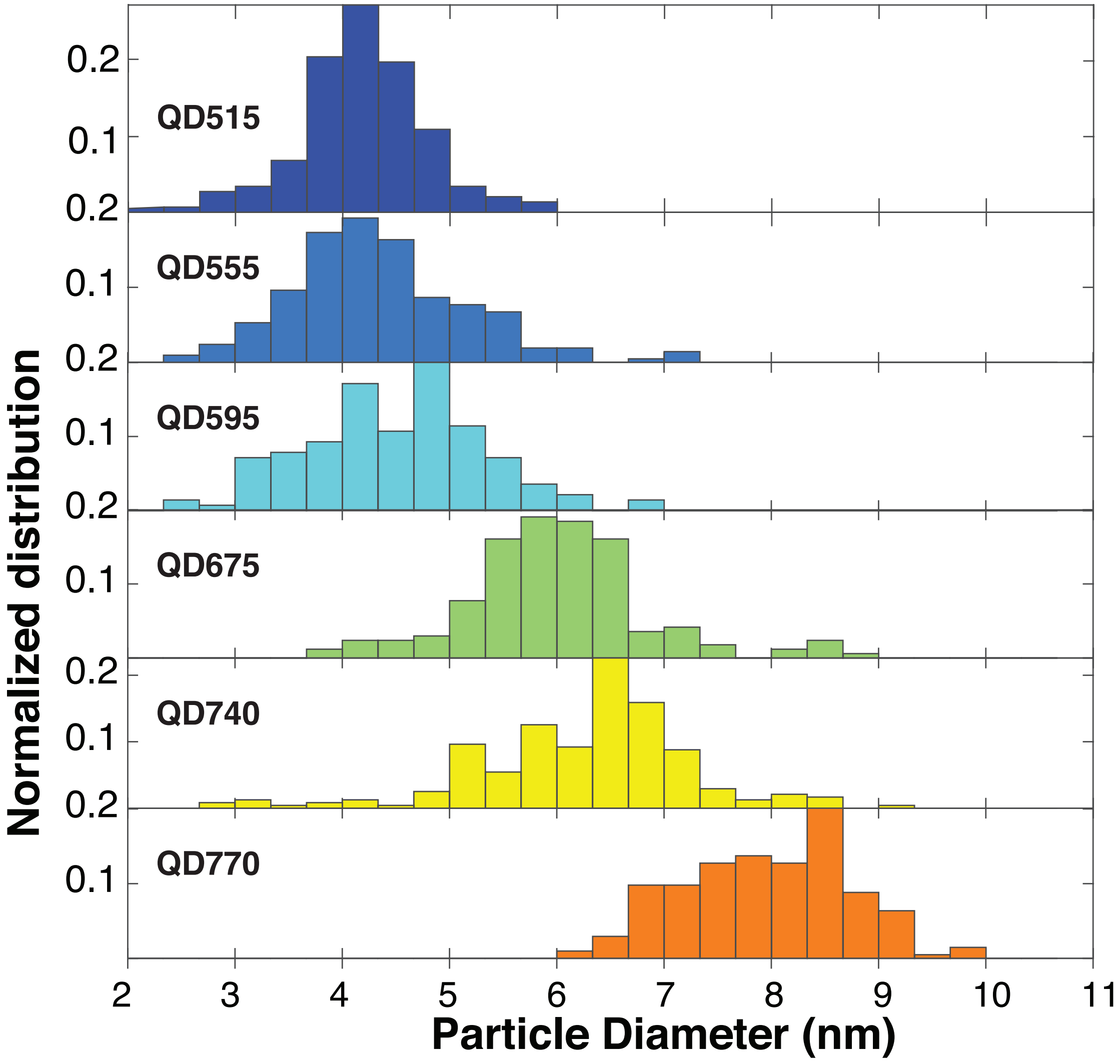


**Figure S6. Size distributions for primary QD size series.** ZnSe/InP/ZnS QDs grown on 2.0 nm diameter ZnSe core; 0.33 nm bin size. From top: QD515 (dark blue), QD555 (blue), QD595 (teal), QD675 (green), QD740 (yellow), and QD770 (orange). The particle sizes were determined from BF STEM images for all samples except QD555, which was sized using TEM images. Particle size determination was performed manually using ImageJ software, n = 129-240.

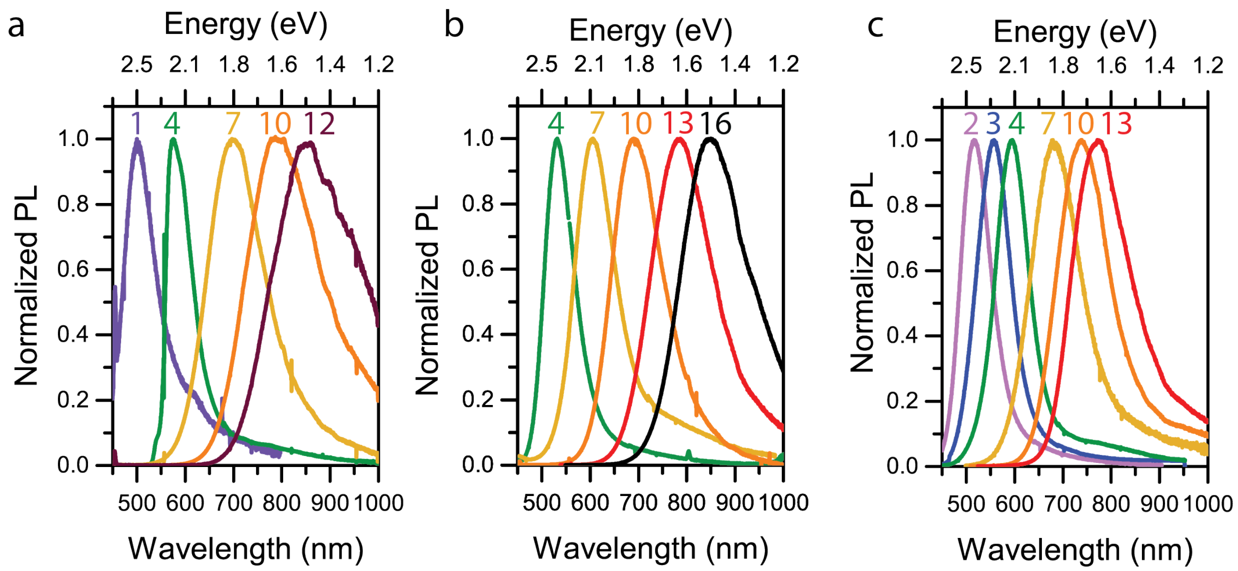


**Figure S7. Repeatability of ZnSe/InP/ZnS synthesis across three reaction series with subtly different reaction conditions.** In each case, sequential PL spectra are shown for samples after the number of InP SILAR rounds indicated as color matched numbers above each corresponding PL spectra. Reaction conditions described in table below.

**Table S1. Summary of QD reactions.**

| Reaction name | Sample name | Reaction conditions | InP SILAR rounds | Emission Peak (nm) |
|---|---|---|---|---|
| QD01-01 | | $D_{ZnSe}$ = 3.26 nm (200 nmol) 10% (v/v) oleylamine in starting solution Samples taken as aliquots from single reaction flask | 1 | 499 |
| QD01-04 | | | 4 | 575 |
| QD01-07 | | | 7 | 697 |
| QD01-10 | | | 10 | 776 |
| QD01-12 | | | 12 | 853 |
| QD02-01 | | $D_{ZnSe}$ = 3.10 nm (200 nmol) 20% (v/v) oleylamine in starting solution Samples taken as aliquots from a single reaction | 1 | 490 |
| QD02-04 | | | 4 | 532 |
| QD02-07 | | | 7 | 599 |
| QD02-10 | | | 10 | 690 |
| QD02-13 | | | 13 | 777 |
| QD02-16 | QD845 | | 16 | 845 |
| QD03-02 | QD515 | $D_{ZnSe}$ = 2.00 nm (100 nmol) 20% (v/v) oleylamine in starting solution Samples are identical reactions taken to different shell thicknesses in parallel flasks | 2 | 515 |
| QD03-03 | QD555 | | 3 | 555 |
| QD03-04 | QD595 | | 4 | 595 |
| QD03-07 | QD675 | | 7 | 675 |
| QD03-10 | QD720 | | 10 | 720 |
| QD03-13 | QD770 | | 13 | 770 |

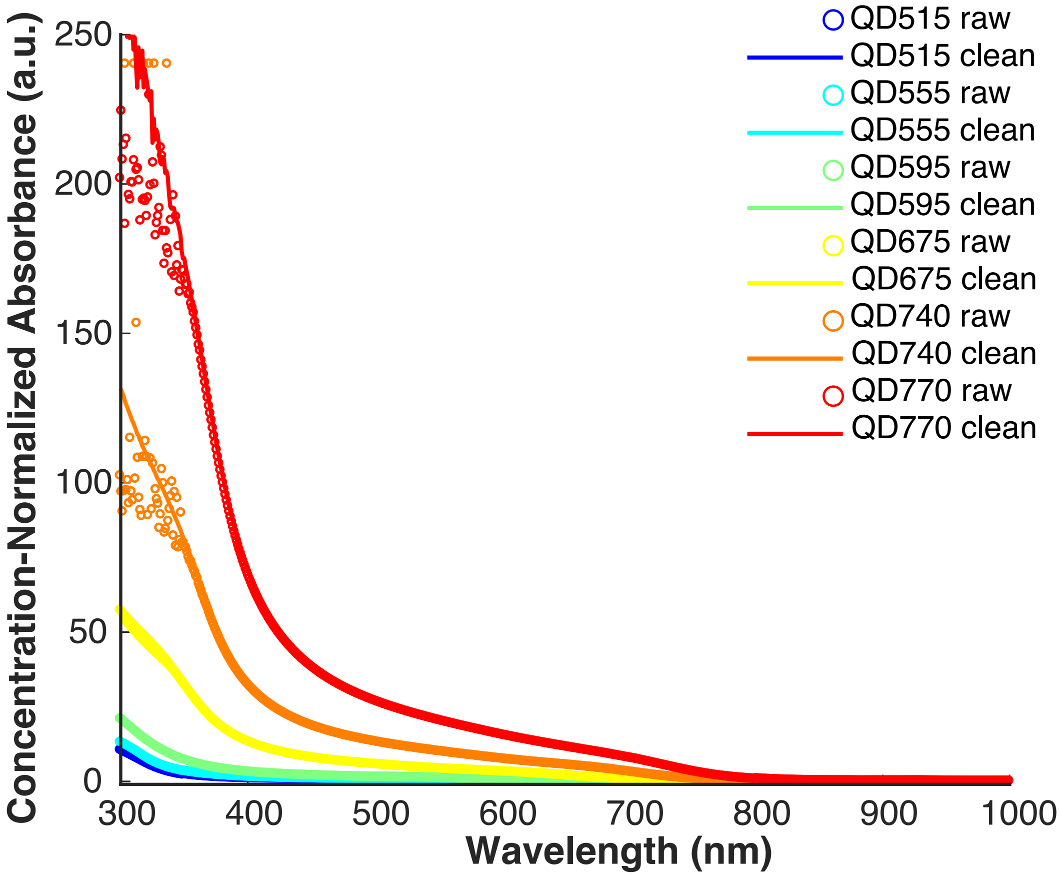


**Figure S8**. **Concentration-normalized absorbance spectra for the QDs.** The circles represent the absorbance of the raw reaction solution after completion diluted in hexanes, with a known concentration of QDs. The lines represent the spectra of cleaned QDs, multiplied by a scaling factor to match the raw solutions at the 1S peak.

**Table S2. Absolute QY measurements for QDs after different treatments.**

| Sample | Cleaned QY (%)[a] | Highest TOP Treated QY Measured (%)[a,b] [Emission Peak (nm)] | QYs of Micelle Encapsulated particles (%)[a,c] | |
|---|---|---|---|---|
| | | | before | after |
| **QD515** | 22.7 ± 0.3 | | | |
| **QD555** | 27.9 ± 0.5 | | | |
| **QD595** | 25.3 ± 0.3 | 36.6 ± 0.4 [560] | | |
| **QD675** | 18.1 ± 1.4 | 38.2 ± 0.1 [673] | 22.9 ± 0.3 | 15.9 ±0.1 |
| **QD740** | 7.6 ± 0.3 | 36.2 ± 0.2 [720] | 22.8 ± 0.2 | 17.8 ± 0.1 |
| **QD770** | 2.7 ± 0.2 | 16.6 ± 0.1 [758] | | |
| **QD845** | 4.0 ± 0.3 | 13.7 ± 0.1 [793] | 13.7 ± 0.1 | 11.9 ± 0.1 |
| **QDot 705 ITK™ [d] (commercial)** | 34.8 ± 0.6 | | 34.8 ± 0.6 | 15.2 ± 0.3 |

[a] The Nanolog® Spectrofluorometer with a Synapse CCD camera and iHR320 fully automated spectrometer (HORIBA Jobin Yvon) is calibrated to measure absolute QYs from 300 – 800 nm. In order to estimate the correct quantum yield for emission > 800 nm, integration of the sample emission spectra was used to determine what fraction of the emitted photons were not included in the QY measurement, and the QY was then adjusted accordingly.
[b] TOP treatment can cause blue shifting of the emission peak, so the peak position of the post-treatment emission is noted in brackets.
[c] The QYs of the specific samples used for *in vivo* imaging both immediately before and after micelle encapsulation.
[d] For comparison, commercially available NIR-emitting QD705s were purchased from ThermoFisher. As shipped (in decane), these QDs exhibited 44.5 ± 0.1% QY. The "before" QY reported is after the QDs were precipitated and resuspended for the encapsulation procedure.

**Table S3. Photoluminescence lifetime tri-exponential fit parameters.**

| | $\tau_1$ (ns) | $A_1$ (%) | $\tau_2$ (ns) | $A_2$ (%) | $\tau_3$ (ns) | $A_3$ (%) | $\bar{\tau}$ (ns) |
|---|---|---|---|---|---|---|---|
| **QD515** | 11.3 | 6.32 | 45.9 | 59.6 | 122 | 34.1 | 91.2 |
| **QD555** | 6.98 | 2.81 | 39.3 | 49.5 | 95.6 | 47.7 | 78.6 |
| **QD595** | 9.15 | 10.1 | 36.0 | 59.0 | 96.1 | 30.9 | 69.9 |
| **QD675** | 10.6 | 5.52 | 51.8 | 45.7 | 142 | 48.8 | 118 |
| **OS740** | 7.49 | 7.62 | 38.7 | 42.5 | 116 | 49.9 | 98.5 |
| **QD770** | 7.84 | 5.67 | 40.5 | 39.1 | 120 | 55.2 | 104 |
| **QD845** | 12.7 | 5.37 | 54.9 | 33.4 | 185 | 61.2 | 166 |

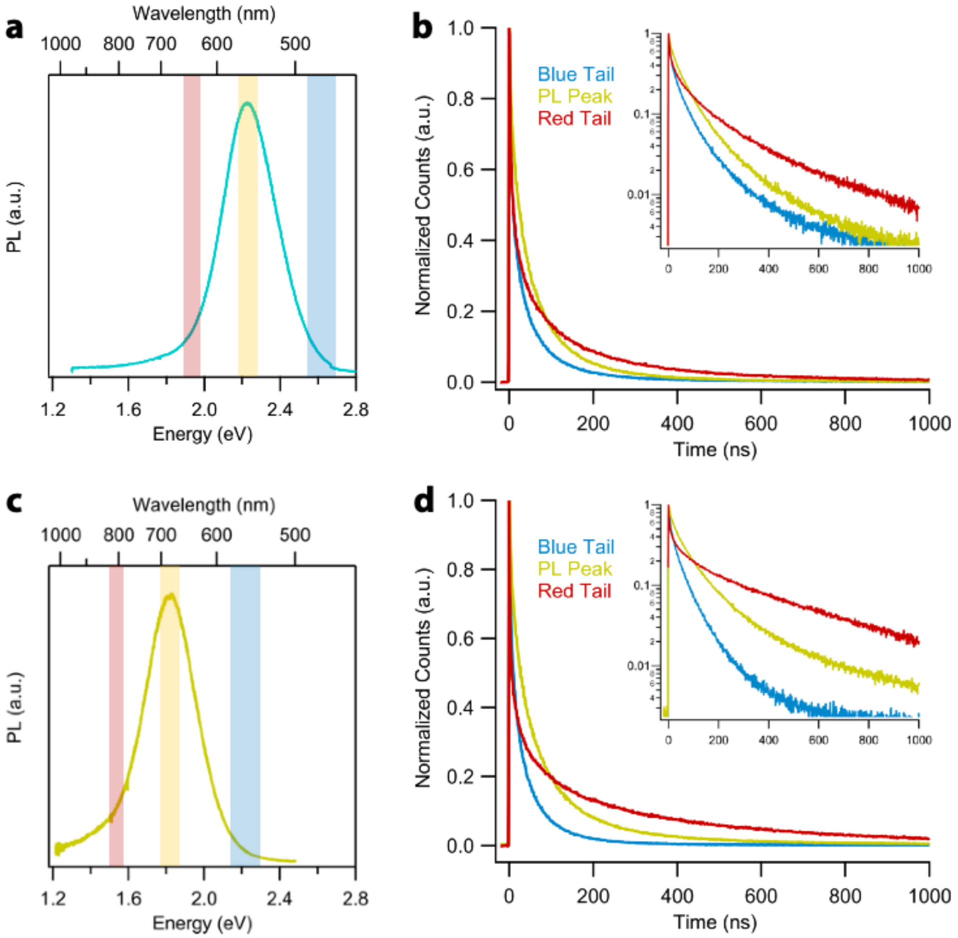


**Figure S9. Emission wavelength-resolved photoluminescence lifetime decay curves** for (**a**,**b**) QD555 and (**c**,**d**) QD675. The shaded regions in (**a**,**c**) illustrate the range over which the lifetimes presented in (**b**,**d**) were measured.

**Table S4. Photoluminescence lifetime tri-exponential fit parameters for the blue tail, peak, and red tail regions of the emission spectra for QD555 and QD675 shown in Figure S11.**

| | $\tau_1$ (ns) | $A_1$ (%) | $\tau_2$ (ns) | $A_2$ (%) | $\tau_3$ (ns) | $A_3$ (%) | $\bar{\tau}$ (ns) |
|---|---|---|---|---|---|---|---|
| **QD555 (blue tail)** | 9.2 | 9.6 | 47.8 | 55.5 | 160.6 | 35.0 | 83.6 |
| **QD555 (peak)** | 16.0 | 8.1 | 62.4 | 59.1 | 194.2 | 32.8 | 101.9 |
| **QD555 (red tail)** | 12.9 | 5.8 | 89.6 | 37.3 | 333.0 | 56.9 | 223.6 |
| **QD675 (blue tail)** | 7.7 | 7.9 | 38.7 | 53.4 | 103.4 | 38.7 | 61.3 |
| **QD675 (peak)** | 22.4 | 10.0 | 84.5 | 58.1 | 300.0 | 31.9 | 147.1 |
| **QD675 (red tail)** | 15.8 | 2.9 | 102.3 | 17.4 | 512.0 | 79.7 | 426.3 |

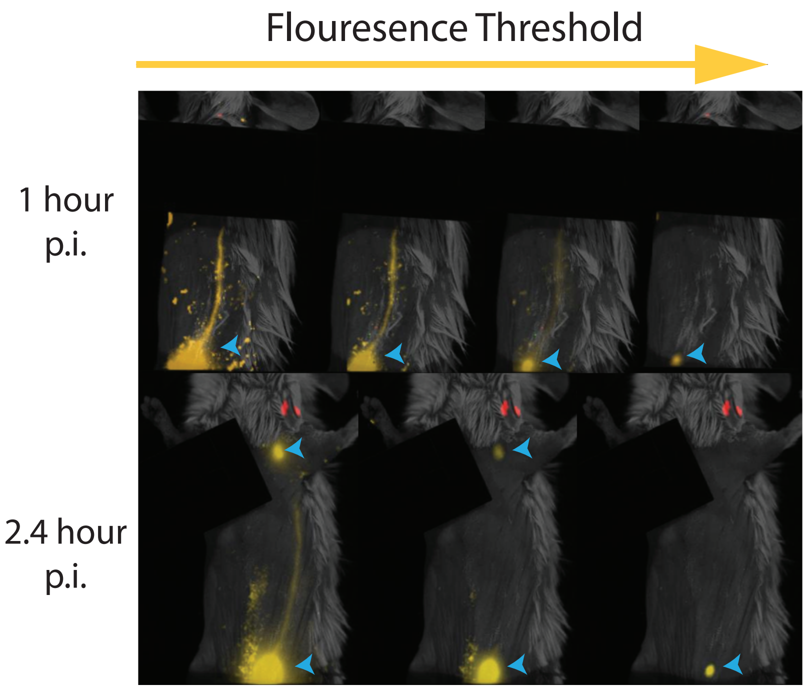


**Figure S10. Using image thresholding to visualize QD drainage in the lymphatic system.** Unmixed fluorescence images at 1 hr and 2.4 hr p.i. of the left side of a mouse injected with a hock injection in the left leg (as well as another hock injection in the right leg and an armpit injection under right forelimb, hidden with an isolator to allow for longer exposure times). The threshold is gradually increased to allow for the visualization of QD lymphatic vessel drainage as well as QD accumulation in the inguinal node in the 1 hr images. For the 2.4 hr p.i. images, the thresholding is adjusted to show QD drainage through the collecting lymphatic vessel, left axillary node, and left inguinal node.

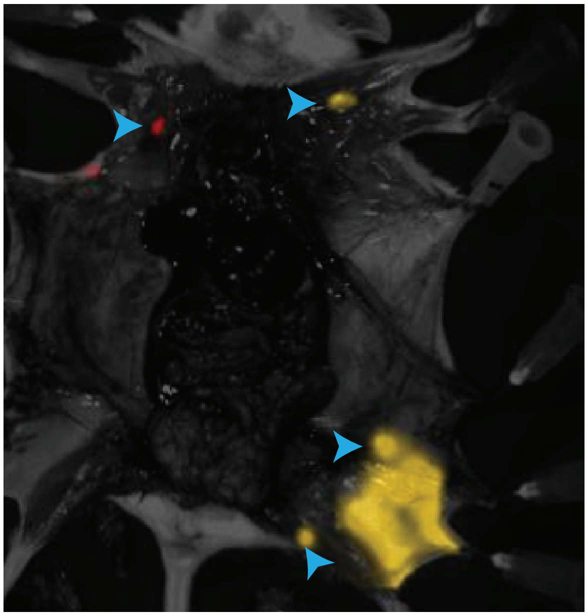

**Figure S11.** Image of the flayed mouse showing localization of the QDs to LNs at 52 hr. p.i. Images taken on Perkin Elmer IVIS Spectrum and analyzed using Living Image software. Blue arrowheads indicate LNs.
